# Functional connectivity strength within the auditory forebrain is altered by song learning and predicts song stereotypy in developing male zebra finches

**DOI:** 10.1101/657825

**Authors:** Elliot A. Layden, Kathryn E. Schertz, Marc G. Berman, Sarah E. London

## Abstract

Much as humans acquire speech in early childhood, the zebra finch (*Taeniopygia guttata*) songbird learns to sing from an adult “tutor” during the first three months of life. Within a well-defined critical period (CP), juvenile zebra finches memorize a tutor song that will guide subsequent motor patterning. This sensory learning process is mediated by tutor experience-dependent neuroplasticity within the auditory forebrain. Here, we used longitudinal resting-state fMRI analyses to investigate whether tutor experience also modifies patterns of functional connectivity (FC) within the juvenile zebra finch brain. Eighteen male zebra finches (only males sing) were scanned before, during, and at the end of the CP, as well as at the young adult stage. Prior to the onset of the CP, birds were separated into rearing conditions: Normal (aviary-housed; *N*=5), Tutored (one adult male tutor and one adult female; *N*=7), and Isolate (two adult females, isolated from male song; *N*=6). Brain-wide voxel-wise analyses identified a single cluster overlapping the left caudomedial nidopallium (NCM) of the auditory forebrain that showed developmentally decreasing FC strength in Isolates but stable or increasing FC in Normal and Tutored birds. Additionally, FC between left NCM and left dorsal cerebellum showed a parallel developmental difference. Developmental changes in left NCM FC strength statistically mediated condition-related differences in song stereotypy. These results extend previous reports of tutor experience-dependent plasticity in NCM at epigenetic, genomic, molecular, and cellular levels to the whole-brain functional network level by demonstrating that tutor experience also influences the development of NCM FC. Moreover, these results link NCM FC to the emergence of song stereotypy.

## Introduction

What neural mechanisms support the learning of complex behaviors during development? The zebra finch (*Taeniopygia guttata*) songbird is a tractable model organism in which to study this fundamental question (Fee & Scharff, 2010; London, 2017). Much as humans acquire speech in early childhood, male zebra finches (females cannot sing) learn to sing from an adult male “tutor” within the first three months of life (London, 2017; Tchernichovski, Mitra, Lints, & Nottebohm, 2001). The foundation of learned song is the formation of a tutor song memory that serves as a “template” to guide subsequent motor patterning. Tutor song memorization occurs within a single restricted developmental phase, or critical period (CP), during which tutor experience has profound and persistent effects on the brain and behavior. The “open” of the CP for tutor song memorization, occurring around post-hatch (P) day 30 (Eales, 1985, 1987; Roper & Zann, 2006), likely depends on neural maturation events that enable learning to commence (London, 2017). In contrast, the CP “close,” after which a zebra finch male can no longer memorize additional songs, is believed to result from tutor experience-dependent neural plasticity remodeling the underlying neural circuitry sufficiently to shift the balance from plasticity to stability. Accordingly, if a juvenile zebra finch is isolated from tutor experience during the CP (“Isolates”), learning potential remains high (Eales, 1985, 1987; Jones, ten Cate, & Slater, 1996, 1992; Morrison & Nottebohm, 1993) well after the typical closure of the CP at P65 (Böhner, 1990; Braaten, 2010; Clayton, 1987; Eales, 1985, 1987; Morrison & Nottebohm, 1993; Roper & Zann, 2006; Slater, Richards, & Mann, 1991). This insight presents the unique opportunity to disambiguate the neural effects of experience-dependent plasticity from the programmed outcomes associated with maturational age (Fee & Scharff, 2010; London, 2017). Moreover, the ability to control tutor experience, and thereby manipulate the closing of the CP, may permit the discovery of neural mechanisms that more broadly promote or limit the ability to learn.

The association regions of the zebra finch auditory forebrain, the caudomedial nidopallium (NCM) and caudomedial mesopallium (CMM), are necessary for the first stage of song learning, i.e., tutor song memorization. For example, pharmacological disruption of molecular signaling pathways within the juvenile zebra finch auditory forebrain during tutor song exposures results in diminished tutor song copying upon maturation (Ahmadiantehrani & London, 2017; London & Clayton, 2008). Specifically, London and Clayton (2008) demonstrated this via inhibition of the extracellular signal-regulated kinase (ERK) signaling pathway (London & Clayton, 2008), which regulates immediate early gene (IEG) *ZENK* expression within the auditory forebrain (Cheng & Clayton, 2004) and is part of a signaling cascade important for long-term potentiation (Bozon et al., 2003). Similarly, Ahmadiantehrani and London (2017) found that either inhibition or constitutive activation of the mechanistic target of rapamycin (mTOR) signaling cascade, also known to contribute to learned behavior (Garza-Lombó & Gonsebatt, 2016; Giovannini & Lana, 2016; Hoeffer & Klann, 2010), decreased the fidelity of tutor song copying. Moreover, several immunocytochemistry studies have found that the density of IEG induction within NCM in response to tutor song playback positively correlates with the strength of song copying in both awake (Bolhuis, Hetebrij, Boer-Visser, Groot, & Zijlstra, 2001; Bolhuis, Zijlstra, Boer-Visser, & Zee, 2000; Terpstra, Bolhuis, & Boer-Visser, 2004) and sleeping (Gobes, Zandbergen, & Bolhuis, 2010) zebra finches. Finally, electrophysiological evidence indicates that neurons within the auditory forebrain are selectively tuned to tutor song in both juveniles (Yanagihara & Yazaki-Sugiyama, 2016) and adults (Phan, Pytte, & Vicario, 2006). Given this converging evidence, the auditory association regions NCM and CMM may present opportune targets in which to examine the neural mechanisms that underlie the CP for tutor song memorization (London, 2017).

While it is known that *local* molecular signaling within the auditory forebrain is essential for tutor song memorization (Ahmadiantehrani & London, 2017; London & Clayton, 2008), much less is known regarding the functional network correlates of tutor experience. That is, how does tutor experience shape functional interactions between the auditory forebrain and the rest of the zebra finch brain, potentially including components of the traditional zebra finch song network (Akutagawa & Konishi, 2010; Bauer et al., 2008; Fortune & Margoliash, 1995; Shaevitz & Theunissen, 2007; Vates, Broome, Mello, & Nottebohm, 1996)? Here, we sought to inform this question by leveraging the advantages of longitudinal resting-state functional MRI (rs-fMRI) functional connectivity (FC) analyses. This methodological paradigm allows for non-invasive, whole-brain, longitudinal data collection, which may be particularly suitable for characterizing a developmental process such as song learning. Moreover, we have previously demonstrated the validity and efficacy of this paradigm in zebra finches (Layden, Schertz, London, & Berman, 2019). Here, we quantified resting-state FC as the temporal correlations embedded within infra-slow (< 0.1 Hz) intrinsic brain activity, recorded as blood-oxygen-level-dependent (BOLD) signals (Biswal et al., 2010; Biswal, Yetkin, Haughton, & Hyde, 1995; Mitra et al., 2018). We then analyzed the voxel-wise developmental trajectory of the intrinsic connectivity contrast (ICC; Martuzzi et al., 2011), a measure that summarizes the centrality or importance of a given voxel or brain region within large-scale FC networks. To interrogate which specific functional connections underlie the ICC levels observed, we conducted seed-to-voxel (S2V) analyses, in which FC is quantified between a cluster of voxels that serve as a “seed” region and voxels distributed throughout the rest of the brain.

We scanned eighteen male zebra finches at four ages: before (P25), during (P45), and at the end of the CP (P65) for tutor song memorization, as well as at the young adult stage (P90). Prior to the onset of the CP, birds were assigned to one of three rearing conditions. *Tutored* birds were raised with one adult male tutor and one adult female non-tutor, whereas *Isolate* birds were raised with two adult females, who do not sing. Thus, a comparison between the Tutored and Isolate conditions allowed us to directly test for effects arising from tutor experience, while holding constant the level of social complexity (i.e., the same number of conspecifics in each condition). Additionally, we included a *Normal* condition, in which birds were raised in an aviary with access to multiple potential adult male tutors, more closely mimicking conditions found in the wild. A contrast between the Normal and Tutored conditions therefore allowed us to gauge any FC effects related to the complexity of social interactions, while holding constant song tutoring. Given the auditory forebrain’s required role in tutor song memorization, we hypothesized that Isolates would exhibit an altered pattern of ICC during ontogeny within the auditory forebrain across the CP compared to either Tutored or Normal birds. However, to conduct a more stringent test of this hypothesis, we conducted data-driven whole-brain analyses that would also allow for the detection of any rearing condition-related developmental differences in ICC, even outside of these *a priori* hypothesized regions, including regions not currently known to be involved in song learning.

While our primary goal was to identify FC signatures of song tutoring, we also sought to link any FC signatures identified to a behavioral outcome, song stereotypy. Although tutor song memorization provides the initial foundation for song learning, a partially overlapping sensorimotor learning phase refines and stabilizes song structure in tutored males (Arnold, 1975; Tchernichovski et al., 2001). Song isolation during the CP (P30-65) results in increased song variability, likely because there is no tutor memory to guide structure (Eales, 1987; Jones et al., 1996), although Isolate songs may become somewhat stereotyped by P100-120 (Morrison & Nottebohm, 1993). Given these findings, we predicted that non-tutored birds in our sample would exhibit lower stereotypy than tutored birds at P65, during the plastic song phase, and potentially at P90 as well, toward the end of the song crystallization phase. Further, if we successfully identified FC signatures of song tutoring, we sought to gauge whether these FC signatures also predicted and/or mediated any differences in stereotypy levels between tutored and non-tutored birds. Our prior hypothesis regarding altered ICC within the auditory forebrain is compatible with this possibility, given a number of indications that auditory forebrain connectivity may play an important role in successful sensorimotor learning (Akutagawa & Konishi, 2010; Bauer et al., 2008; Fortune & Margoliash, 1995; Shaevitz & Theunissen, 2007; Vates et al., 1996; Yanagihara & Yazaki-Sugiyama, 2016).

## Method

### Animals

All animal procedures were approved by the Institutional Animal Care and Use Committee of the University of Chicago in accordance with the NIH Guide for the Care and Use of Laboratory Animals. All birds were maintained under a 14/10 hour light/dark photoperiod throughout the experiment, with food and water provided *ad libidum*. We collected rs-fMRI scans for male zebra finches at four ages, capturing FC at various stages of the song learning process: prior to the CP for tutor song memorization (P25), in the approximate middle of the CP (P45), at the end of the CP (P65), and toward the end of the sensorimotor rehearsal period, when tutor song copying is complete and song stereotypy is high (P90). Zebra finches were assigned to one of three rearing conditions, sorted to separate brothers and to temporally batch birds into all of the rearing conditions following their P25 rs-fMRI scanning session, and before the onset of the CP at around P30 (Roper & Zann, 2006): *Normal* (aviary housed with access to multiple potential adult male tutors; *N*=5), *Tutored* (housed with one adult male tutor and one adult female; *N*=7), and *Isolate* (housed with two adult females, meaning no access to song tutors but having experience with conspecific vocalizations (females produce innate calls), and with the same complexity of social interactions as the Tutored group; *N*=6). Importantly, housing Isolate birds with two adult females allowed us to measure effects specific to isolation from song, as opposed to social isolation more generally.

One Normal male died of unknown cause in the group aviary between P65 and P90 (unrelated to scanning). Additionally, a P25 scan was not successfully obtained for two Tutored birds and one Isolate bird, due to a scanner coil hardware failure. This yielded a total of 68 out of 72 possible successful scanning sessions across 18 male zebra finches. While the original head coil was sent out for repairs, a substitute head coil was used to scan seven P25 birds (2 Normal, 2 Tutored, and 3 Isolate), whereas the repaired original was used to scan all other target ages. To statistically control for any potential effects resulting from the use of different head coils, we included a categorical covariate denoting which head coil was used for scanning in all of the following analyses that utilized P25 scans. Importantly, this issue only affected our P25 baseline data and not our target ages of primary interest (ages P45-P90).

Finally, due to weekday-only scanner availability and scheduling conflicts, not all birds could be scanned on exact target days, but the age range was kept as close as possible to the targets: P25 (range: P24-P26), P45 (range: P44-P46), P65 (range: P63-P67), and P90 (range: P88-P91). Importantly, age at scan time did not significantly vary by condition at any target age, nor across target ages (ANOVA, all *p*’s > 0.17). Additionally, precise ages were recorded for each bird, and age was implemented as a continuous variable in all analyses described below.

### Scanning Procedure and Preprocessing

MRI data collection was conducted at the MRIS Facility of the University of Chicago. Upon arrival, zebra finches were anesthetized using an admixture of oxygen and isoflurane gas (1.5-2.25%). Subsequently, a light maintenance dosage of isoflurane was maintained throughout the experiment (0.5-2%). The use of isoflurane is well-established in task-based fMRI studies of zebra finches (Boumans, Theunissen, Poirier, & Van Der Linden, 2007; Poirier, Boumans, Verhoye, Balthazart, & Van der Linden, 2009), and isoflurane was also used successfully in our previous resting-state fMRI zebra finch study (Layden et al., 2019). Importantly, although hemodynamic response magnitude is reduced by isoflurane anesthesia, hemodynamic response shape remains largely unchanged (Zhao, Jin, Wang, & Kim, 2007). Following anesthesia administration, birds were fitted with a temperature probe and respiration monitoring pad, allowing for body temperature, respiratory rate, and isoflurane percentage to be monitored throughout the scanning period. The zebra finches were wrapped in a felt cloth to help maintain body temperature. Upon insertion into the scanner, a warm air feedback system was used to further maintain body temperature within a normal physiological range (40.0 ± 0.2 °C).

#### Imaging Data Acquisition

Neuroimaging data were acquired using a 30 cm bore 9.4 T Bruker small animal MRI scanner. A TurboRARE-T2 Multislice anatomical scan was acquired first during each scanning session (TR = 3.5 s, TE = 20 ms, Matrix Size: 256 x 256, in-plane resolution = 70.3 μm x 70.3 μm, slice-thickness = 200 μm, 59 slices, 9 averages). Resting-state RARE spin-echo T2-weighted MR images were then acquired (TR = 3.2 s, TE_effective_ = 27 ms, 3 echoes per volume, Matrix Size: 128 x 36, in-plane resolution = 141 μm x 500 μm, slice-thickness = 750 μm, 15 slices). Slices were acquired in an interleaved, ascending order. 180 volumes were acquired consecutively with an effective sampling rate of 3.2 seconds per volume, yielding a total resting-state scan time of 9 minutes and 36 seconds. A spin-echo pulse sequence was used due to observations that gradient-echo imaging may be particularly vulnerable to susceptibility artifacts in zebra finch whole-brain fMRI (Poirier & Van der Linden, 2011). To avoid T1-equilibration effects, the first five volumes of each functional series were discarded.

#### Preprocessing

Image preprocessing was completed using a combination of ANTs (Avants et al., 2011; Avants, Tustison, & Song, 2009), AFNI (Cox, 1996), and custom Matlab scripts. First, magnetic field intensity inhomogeneity was corrected for both functional and anatomical images using the “N4” bias field correction algorithm (Tustison et al., 2010). Second, an average of the bias-corrected anatomical scans was used to initialize SyGN (Avants et al., 2010) in ANTs. ANTs registration accuracy compares favorably among commonly used nonlinear deformation algorithms (Avants et al., 2011; Klein et al., 2009). Third, functional scans were corrected for slice timing differences using ANTs ImageMath SliceTimingCorrection. Slice timing correction may be particularly essential in the case of longer duration TRs (Sladky et al., 2011), and it is recommended that this step be performed prior to realignment for interleaved slice acquisitions (Johnstone et al., 2006). Fourth, motion correction was performed using the antsMotionCorr function in ANTs, and the six rigid-body motion parameters were retained for later nuisance regression. Fifth, the average motion-corrected functional scans were affine coregistered to their corresponding bias-corrected anatomical volumes using the antsRegistrationSyN function. Sixth, to minimize interpolations, we applied all transformations obtained from (1) motion correction, (2) coregistration, and (3) structural normalization to the slice-timing corrected functional scans in a single step using the ANTs function antsApplyTransforms. Finally, we used the AFNI function 3dBlurInMask to smooth the functional data within our custom brain mask using a 0.8438 mm FWHM Gaussian kernel (corresponding to 6.00 x 1.69 x 1.12 voxels in the *x*, *y*, and *z* dimensions, respectively). This function avoids contaminating signals at the edge of the brain with signal from other surrounding tissues (or zeros). All subsequent analyses were carried out using normalized data at the functional image resolution (0.141 x 0.500 x 0.750 mm, 8,720 in-mask voxels), and results were interpolated to template resolution for display purposes.

#### Data quality assessment

To assess the quality and consistency of spatial normalization, we computed voxel-wise Pearson correlations between the custom template and normalized anatomical scans across voxels (*r_mean_* = 0.94, *SD* = 0.01). We also performed this assessment for the normalized functional scans, averaged across time (*r_mean_* = 0.86, *SD* = 0.03). Both assessments indicated a robust and consistent spatial normalization across scans, comparable to prior literature (Layden et al., 2017; Layden, Schertz, London, & Berman, 2019). We also assessed the temporal signal-to-noise ratio (*tSNR*) of the functional images and noted robust *tSNR* across scans (*M* = 56.36, *SD* = 13.10). *tSNR* was comparable to other recent fMRI studies utilizing sub-millimeter voxel sizes and high magnetic field strengths (Chang et al., 2018; Liska, Galbusera, Schwarz, & Gozzi, 2015; Yoshida et al., 2016). For additional details on *tSNR* assessment, see Layden et al. (2019).

### Data Extraction and Denoising

We implemented denoising procedures using MRIqual and custom Matlab scripts. Physiological variables (body temperature and isoflurane dose), recorded at one-minute intervals, were interpolated using a clamped cubic spline function to the temporal resolution of our functional series. Then, a nuisance regression was performed in which the six rigid body motion parameters (Bright & Murphy, 2015), body temperature, isoflurane dose, and linear and quadratic trends (Tanabe, Miller, Tregellas, Freedman, & Meyer, 2002) were removed from the voxel-wise time series. Following nuisance regression (Hallquist, Hwang, & Luna, 2013), the residual ROI time series were bandpass filtered (range: 0.008 to 0.1 Hz).

### Intrinsic Connectivity Contrast (ICC) Analyses

We used a data-driven approach to investigate associations between rearing condition and brain-wide FC strength, measured using the ICC, which does not require arbitrary thresholding of functional connections (Martuzzi et al., 2011). ICC is computed at each brain voxel by averaging the squared Pearson correlations between a given voxel time series and every other voxel time series in the brain (Martuzzi et al., 2011); the voxel-wise distribution of ICC is then standardized (i.e., *Z*-scored) within each functional scan (Layden et al., 2017). ICC has previously proven effective for identifying functional hubs within the human brain associated with a variety of affective, social, and cognitive phenomena (e.g., Layden et al., 2017; Moreno-López, Sahakian, Manktelow, Menon, & Stamatakis, 2016; Vatansever, Manktelow, Sahakian, Menon, & Stamatakis, 2017).

Using an LME random intercept model, we regressed voxel-wise ICC onto age, rearing condition, an age by rearing condition interaction, isoflurane dose, body temperature, head coil, and a random intercept for bird. For each voxel, we retained the *t*-statistic summarizing the age by rearing condition interaction effect. Significant clusters of voxels wherein the developmental trajectory of ICC differed by rearing condition were detected using the random field theory-based cluster-extent thresholding method, as implemented in the DPABI toolbox (cluster-defining threshold (CDT): *p-uncorrected* < 0.001, FWE-corrected cluster-extent threshold: *pFWE* < 0.05; Yan et al. 2016).

#### Normal versus Tutored contrast

Although our primary interest was in whether Isolate birds would exhibit a pattern of FC development distinguishable from both Normal and Tutored birds, we first examined whether Normal and Tutored birds were distinguishable via FC. Any such effects could potentially be related to social group size or the number of available tutors, rather than tutor experience or lack thereof.

#### Tutored versus non-tutored contrast

Given that the Normal vs. Tutored contrast did not yield significant results (see Results below), we combined these rearing conditions into a single “Tutored” condition to yield increased statistical power for our contrast of primary interest. We then examined whether the developmental trajectory of ICC differed in any brain regions between tutored (i.e., Normal and Tutored) and non-tutored birds (i.e., Isolates).

### Seed-to-Voxel (S2V) Functional Connectivity Analyses

ICC enables the detection of clusters based on global FC strength but does not reveal the specific functional connections that may underlie any global FC differences observed. We therefore conducted a “seed-to-voxel” (S2V) analysis, in which we extracted the average BOLD signal from the voxels of the significant cluster detected in ICC analyses (see Results). We then computed S2V FC as the Fisher *Z*-transformed Pearson correlation between the cluster/seed signal and the signal of every other brain voxel. At each voxel, we then regressed S2V FC onto age, rearing condition, an age by rearing condition interaction, isoflurane dose, body temperature, head coil, and a random intercept for bird. Finally, clusters of S2V FC showing developmental trajectories that differed by rearing condition were identified using the same cluster-extent thresholding method as for ICC analyses (CDT: *p-uncorrected* < 0.001, cluster-extent threshold: *pFWE* < 0.05).

### Song Stereotypy

Song bouts were recorded in a sound isolation chamber for a period of approximately 24 hours prior to rs-fMRI scans at P65 (*N* = 12) and P90 (*N* = 16). Recording incorporated an overnight period that is known to alter song structure, and excluded songs from the first 3 hours after lights-on because song is more variable first thing in the morning (Derégnaucourt, Mitra, Fehér, Pytte, & Tchernichovski, 2005). Some birds did not sing during the recording period, thus resulting in the different sample sizes for each age.

#### Song stereotypy quantification

We quantified song stereotypy (i.e., self-similarity or consistency across song bouts) using Sound Analysis Pro 2011 (Tchernichovski, Nottebohm, Ho, Pesaran, & Mitra, 2000). Three standard measures of stereotypy were calculated: %-Similarity (a coarse-grained comparison of song segments using 60 ms time windows), Accuracy (a fine-grained comparison of song segments within 15 ms time windows), and %-Sequential (quantifying the extent to which matching sound segments occur with the same ordering in different song bouts). Additional details regarding the calculation of these metrics are available from the Sound Analysis Pro 2011 reference manual (Tchernichovski & Mitra, 2004).

We found that these three measures were highly correlated between birds across ages P65 and P90 (%-Similarity and Accuracy: *r*(26) = 0.89, %-Similarity and %-Sequential: *r*(26) = 0.71, Accuracy and %-Sequential: *r*(26) = 0.78). Moreover, the first principal component (PC) from a principal component analysis conducted on these 3 measures explained 88.2% of total variance among the three stereotypy measures, and this PC loaded (i.e., correlated) highly on each individual measure (loadings: %-Similarity (*r* = 0.94), Accuracy (*r* = 0.96), %-Sequential (*r* = 0.89)). To avoid issues with both multicollinearity and multiple comparisons, we utilized the first PC of the three stereotypy measures as our variable of interest for subsequent analyses, and for simplicity, we will henceforth refer to this first PC as “stereotypy.”

#### Condition-related differences in stereotypy

We predicted that non-tutored birds would exhibit less stereotyped songs than Tutored birds at P65, during the plastic song phase, and potentially at P90 as well, toward the end of the song crystallization phase. Additionally, there are indications that the number of song bouts (i.e., the amount of practice) during the plastic song phase strongly predicts stereotypy at P112, whereas the number of song bouts during the song crystallization phase does not predict stereotypy at P112 (Johnson, Soderstrom, & Whitney, 2002). Therefore, we hypothesized that the four birds (2 Normal, 2 Tutored) who sang at P90 but not at P65 would exhibit lower stereotypy at P90. To test these hypotheses, we implemented an LME model in which stereotypy served as the criterion variable, and the following were included as fixed-effects predictors: rearing condition (tutored vs. non-tutored), age, and a nominal factor denoting whether or not each bird sang at P65; a random intercept was included for bird. With only two ages (P65 and P90) per bird, we found that the Hessian matrix was not positive definite for this LME model (i.e., the random intercept was not supported), so we dropped the random intercept and implemented the model using ordinary least squares (OLS) regression.

#### Predicting song stereotypy using FC features

We next investigated whether the cluster(s) identified based on ICC or S2V FC analyses predicted song stereotypy. To do so, we selected an optimal model for predicting stereotypy from among the FC features using the ‘step’ function from the “lmerTest” package (Kuznetsova, Brockhoff, & Christensen, 2017) in R. We performed stepwise model reduction based on reductions in AIC, requiring significant differences between nested models based on likelihood-ratio tests. The initial model included the following predictors: age, condition, average ICC of the significant cluster identified, developmental change in ICC from the previous age (i.e., P65 ICC minus P45 ICC predicting P65 stereotypy and P90 ICC minus P65 ICC predicting P90 stereotypy), average S2V FC of the significant cluster identified, and developmental change in S2V FC from the previous age. Notably, we controlled each FC feature for any effects of body temperature and isoflurane dose prior to stepwise selection. Additionally, we included the developmental change in ICC and S2V FC across ages because we reasoned that developmental changes might more strongly predict stereotypy than FC measured within a single age. This could be the case if, for example, low ICC during the CP for tutor song memorization promotes brain network flexibility in support of high learning potential, and relative increases in ICC at ages beyond the CP promote brain network stability in support of crystallization. Similarly, it is possible that the magnitude of developmental change in FC is more important than the absolute level of FC, which may vary randomly between birds.

#### FC mediation of stereotypy differences between rearing conditions

If the ICC or S2V FC features described in the previous section proved to be predictive of stereotypy, we planned to empirically test whether they statistically mediated stereotypy differences between rearing conditions (i.e. tutored vs. non-tutored). We used the “mediation” package in R (Tingley, Yamamoto, Hirose, Keele, & Imai, 2014) to estimate the average causal mediation effect (*ACME*), which corresponds to the proportion of the rearing condition effect mediated by a given FC feature. We also estimated the average direct effect (*ADE*), which corresponds to the proportion of the rearing condition effect not mediated by the FC feature. Lastly, we estimated the total effect, which is a combination of the *ACME* and *ADE*. All effects were estimated using the non-parametric bias-corrected and accelerated bootstrap method with 10,000 iterations.

## Results

### Intrinsic Connectivity Contrast (ICC) Analyses

#### Normal versus Tutored comparison

No significant clusters were identified (CDT: *p-uncorrected* < 0.001, cluster-extent threshold: *pFWE* < 0.05).

#### Tutored versus Isolate comparison

Voxel-wise LME models identified a significant cluster of 19 functional space voxels (CDT: *p-uncorrected* < 0.001, cluster-extent threshold: *pFWE* < 0.05) centered within the left auditory forebrain (Figure 1). Specifically, 13 voxels (68.4% of the cluster) overlapped a ventral posterior portion of the left auditory forebrain, likely corresponding to NCM, and 6 voxels (31.6% of the cluster) overlapped an unlabeled region directly adjacent to the medial posterior portion of the left auditory forebrain.

**Figure 1.**
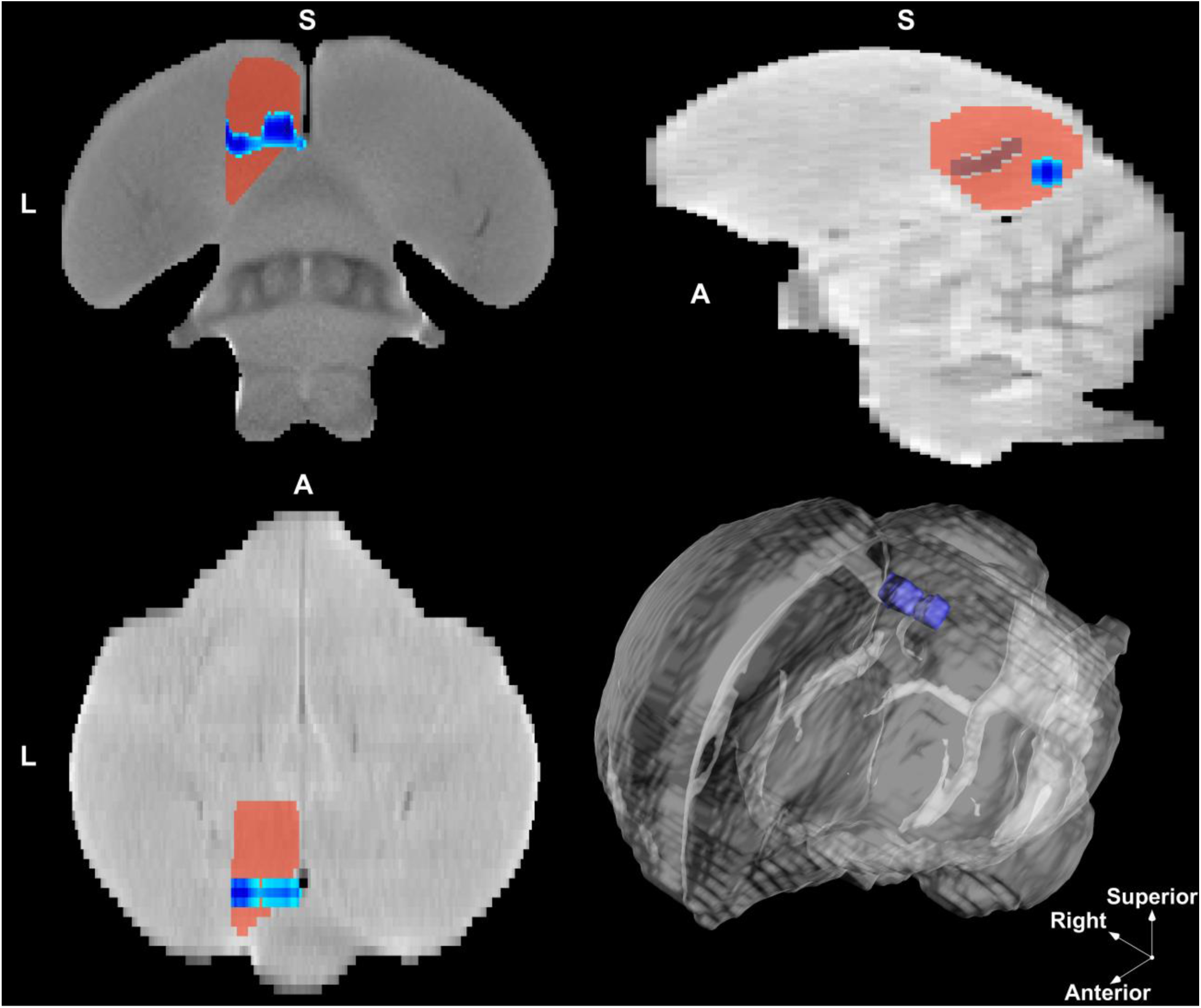
Intrinsic connectivity contrast (ICC) cluster. Non-tutored male zebra finches showed decreasing ICC with age within this cluster, whereas tutored zebra finches showed increasing or stable ICC. From top-left to bottom-right: coronal view (slice 16), sagittal view (slice 148), axial view (slice 169), 3D rendering. All voxels depicted exhibited *t*-statistics ≤ −3.23 (*p* < 0.001) for the rearing condition by age interaction effect.

The average ICC level within this cluster significantly decreased with age in non-tutored relative to tutored birds (*β* = −1.05, *t*(61) = −4.77, *p* < 0.001). A follow-up analysis revealed that the ICC within this cluster marginally *increased* with age for Normal birds (*β* = 0.44, *t*(14) = 1.93, *p* = 0.074), non-significantly *increased* with age for tutored birds (*β* = 0.21, *t*(21) = 1.31, *p* = 0.204), and significantly *decreased* with age for Isolate birds (*β* = −0.65, *t*(18) = −3.06, *p* = 0.007). The developmental trends in ICC are displayed for each individual bird in Figure 2.

**Figure 2.**
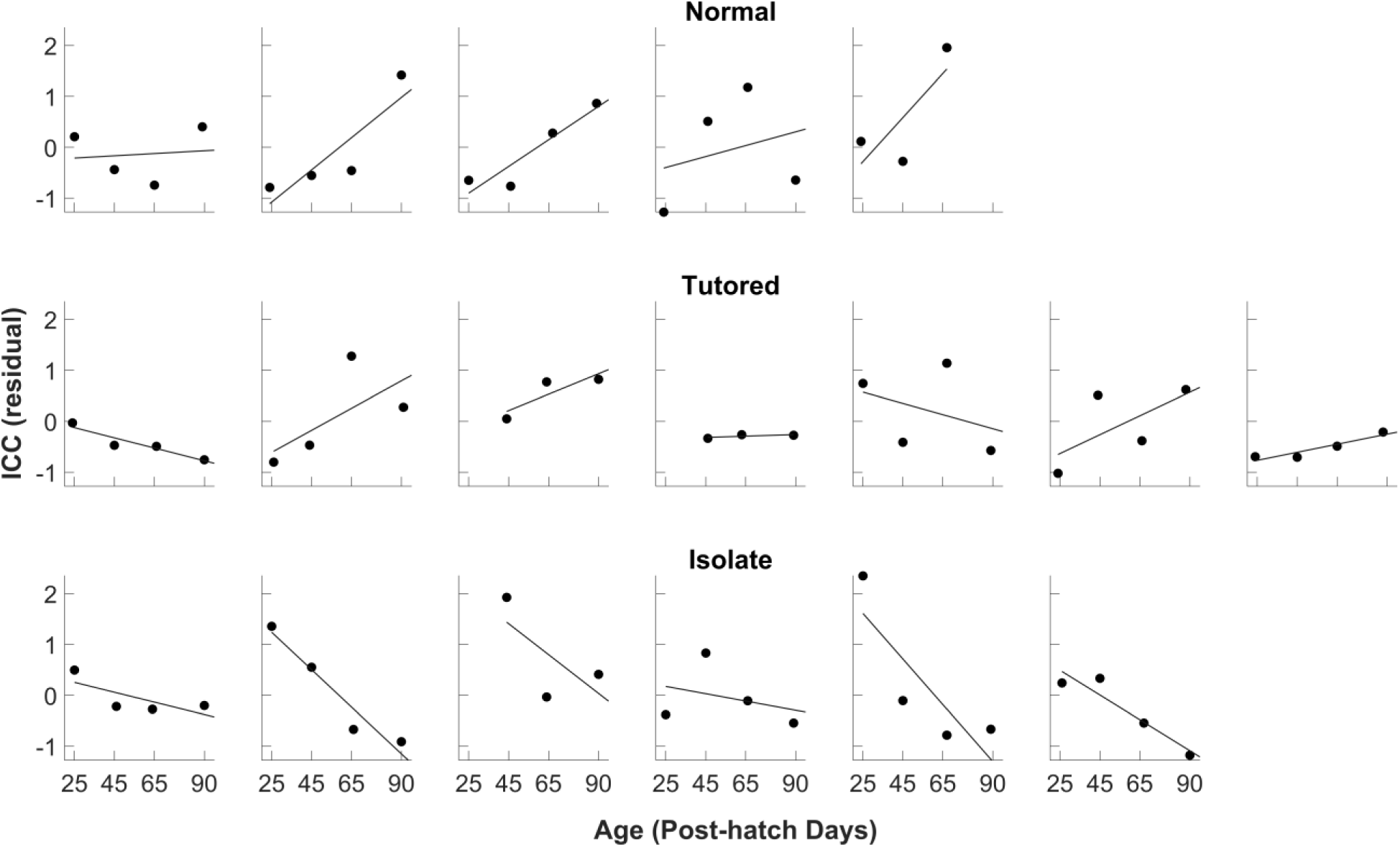
Within-subjects ICC developmental trends for the left NCM cluster. Separate scatter plots show the developmental trajectory of ICC across ages P25 to P90 (*x*-axis) for each individual bird. A within-subjects least-squares trend line is displayed for reference. ICC scores (*y*-axis) are residuals from a regression controlling ICC across birds for the effects of body temperature, isoflurane dose, and head coil.

#### Testing for lateralization

Given that the left NCM cluster appeared to be located exclusively within the left hemisphere, we conducted a follow-up analysis to test whether a similar effect could also be observed in a mirror-symmetric (i.e., homotopic) cluster within the right hemisphere, or if, alternatively, the effect was truly left lateralized. Notably, we previously found that the auditory forebrain exhibited strong bilateral (homotopic) FC (Layden et al., 2019). Here, we found that the average ICC level within a homotopic right-hemispheric cluster also significantly decreased with age in non-tutored relative to tutored birds (*β* = −0.89, *t*(61) = −3.92, *p* = 0.002). This developmental difference was numerically smaller than that observed in the left hemisphere (*β* = −1.05). However, the effect sizes in each hemisphere were not significantly different, as revealed by an LME model in which we tested the three-way interaction between age, rearing condition, and hemisphere as a binary categorical variable (*β* = 0.35, *t*(125) = 1.22, *p* = 0.224). This result indicates that tutor experience may affect the global FC of both left and right NCM, with a numerically but not significantly stronger effect observed in the left hemisphere.

### Seed-to-Voxel (S2V) Analyses

ICC measures the global FC strength of a given voxel or cluster; however, ICC alone does not reveal what specific functional connections underlie the level of global FC observed. To investigate whether region-specific FC also showed a differential developmental trajectory between tutored and non-tutored birds, we used the previously identified left NCM cluster as an FC “seed” region for S2V analyses. We identified one significant cluster of 21 voxels (Figure 3) that showed a rearing condition-related developmental difference in FC to the left NCM cluster seed (CDT: *p-uncorrected* = 0.001, cluster-extent threshold: *pFWE* < 0.05). Two voxels (9.5% of cluster volume) overlapped the left posterior auditory forebrain (i.e., NCM), whereas much of the rest of the cluster appeared to overlap the left dorsal cerebellum.

**Figure 3.**
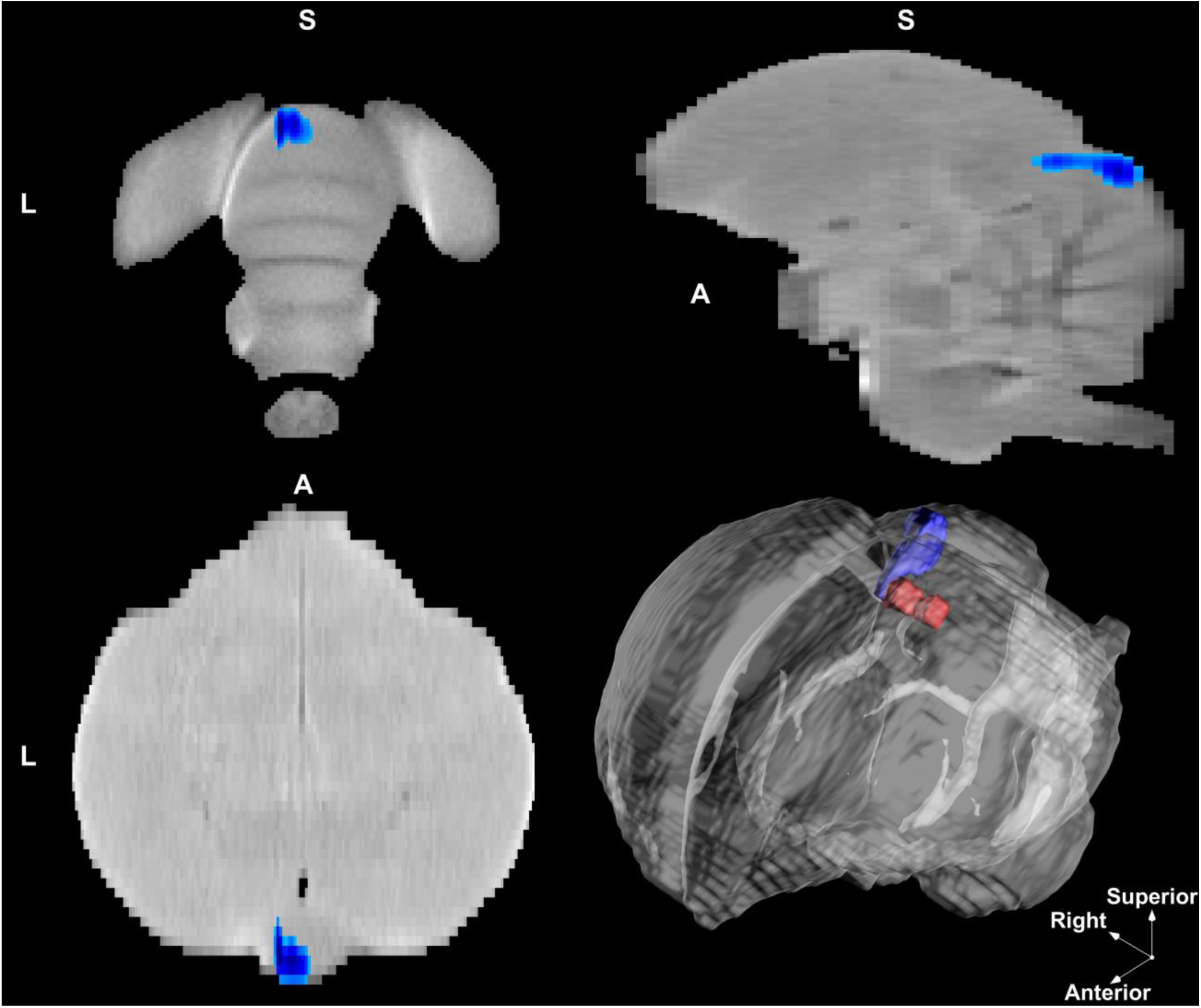
Seed-to-voxel (S2V) FC cluster. The significant cluster of voxels wherein non-tutored male zebra finches, compared to tutored zebra finches, showed developmentally decreasing FC to the previously identified left NCM cluster. From top-left to bottom-right: coronal view (slice 10), sagittal view (slice 145), axial view (slice 172), 3D rendering. All voxels depicted exhibited *t*-statistics ≤ −3.23 (*p* < 0.001) for the rearing condition by age interaction. The red cluster depicted in the 3D rendering (bottom right) is the left NCM cluster from ICC analyses, used here as an FC seed region.

The developmental trajectory of FC for this connection markedly differed between non-tutored and tutored birds (age * non-tutored vs. tutored: *β* = −1.14, *t*(61) = −5.65, *p* < 0.001). Next, we conducted a follow-up analysis in which the age and S2V FC association was measured separately within each rearing condition. In both Normal and Tutored birds, S2V FC non-significantly *increased* with age (Normal: *β* = 0.28, *t*(14) = 1.11, *p* = 0.284; Tutored: *β* = 0.18, *t*(21) = 0.93, *p* = 0.362). In contrast, the S2V FC strongly *decreased* with age in Isolate birds (*β* = −0.94, *t*(18) = −7.15, *p* < 0.001). The developmental trends of S2V FC are shown for each individual bird in Figure 4.

**Figure 4.**
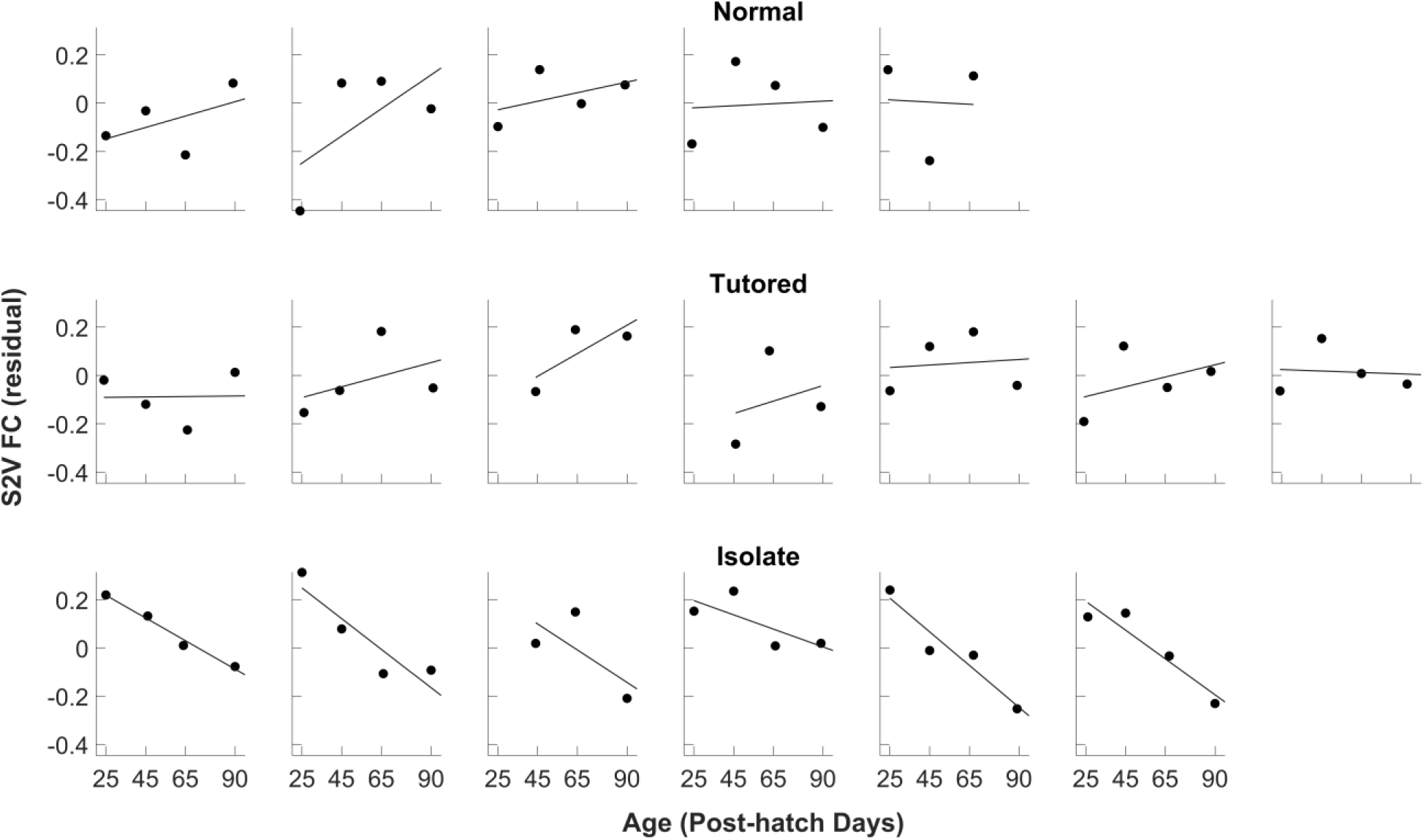
Within-subjects developmental trend of S2V FC. Separate scatter plots show the developmental trajectory of S2V FC across ages P25 to P90 (*x*-axis) for each individual bird. A within-subjects least-squares trend line is displayed for reference. S2V FC values (*y*-axis) are residuals from a regression controlling S2V FC across birds for the effects of body temperature, isoflurane dose, and head coil.

### Song Stereotypy Analyses

We next ran an ordinary least squares (OLS) regression model predicting stereotypy with rearing condition (tutored vs. non-tutored), age, and a factor denoting whether each bird sang at P65. We found that non-tutored birds’ songs were significantly less stereotyped than tutored birds’ across ages P65 and P90 (*β* = −0.97, *t*(24) = −2.69, *p* = 0.013). Additionally, the two Normal and two Tutored birds who sang at P90 but not P65 exhibited markedly less stereotyped songs at P90 than those who sang at P65 (*β* = −1.27, *t*(24) = −2.41, *p* = 0.024). Finally, increasing age from P65 to P90 predicted marginally increased stereotypy (*β* = 0.33, *t*(24) = 1.84, *p* = 0.078). Overall, this model explained 32.6% of the total stereotypy variance (*Adjusted R^2^* = 0.242). The developmental trends in song stereotypy are displayed in Figure 5.

**Figure 5.**
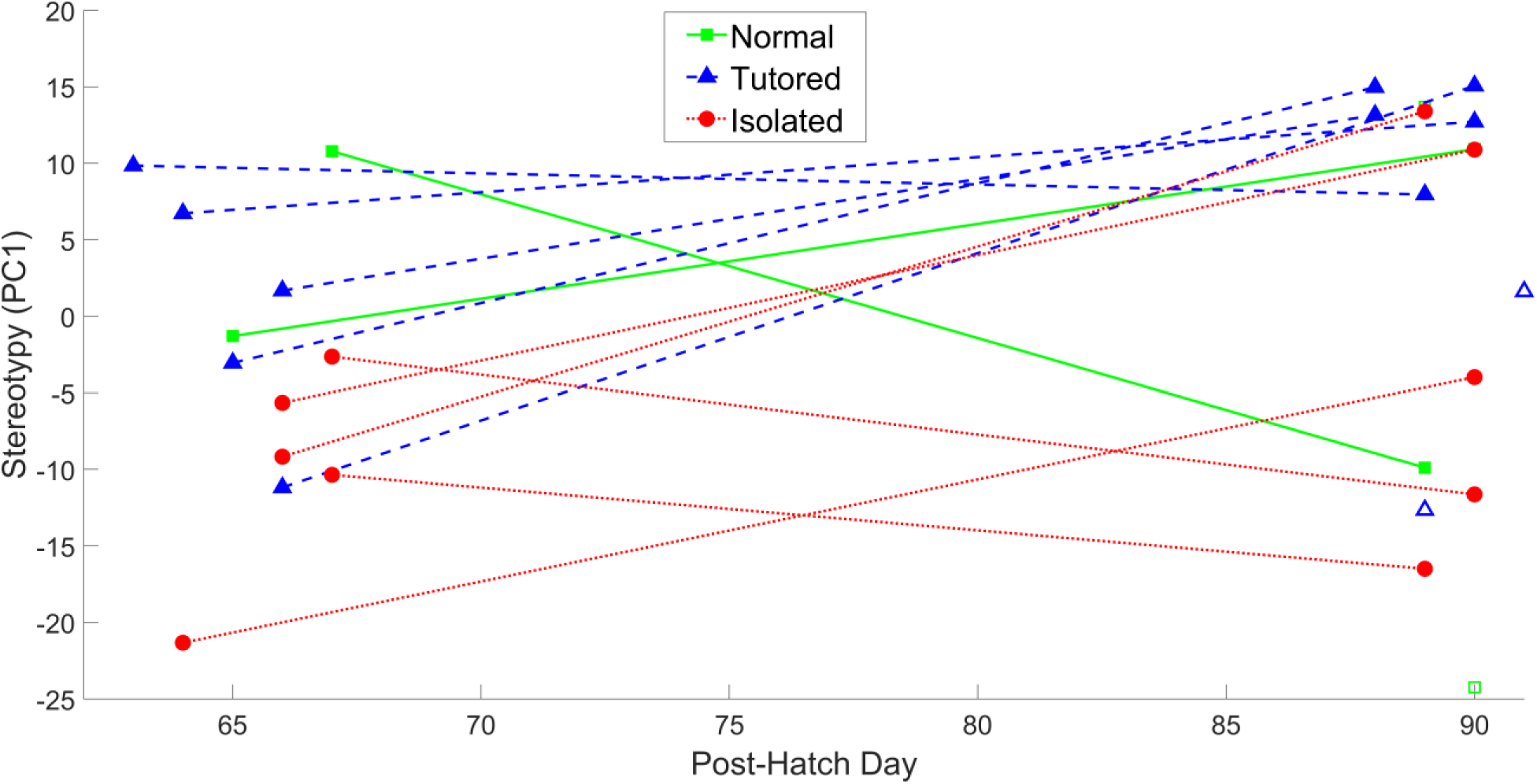
Stereotypy scores by age and condition. Each Normal condition bird is depicted with a solid green line connecting corresponding stereotypy scores at each age (square markers). Tutored birds are depicted with a dashed blue line connecting stereotypy scores (triangle markers). Isolate birds are depicted with red dotted lines connecting stereotypy scores (circle markers). Hollow markers are used to denote the four birds (2 Normal, 2 Tutored) who sang at P90 but not P65. Ages (*x*-axis) correspond to age at rs-fMRI scan date.

A follow-up analysis did not support the inclusion of an age by condition interaction term in our model predicting stereotypy (*F*(1, 23) = 0.04, *p* > 0.80; age by non-tutored: *β* = 0.08, *t*(23) = 0.21, *p* > 0.80). However, separate post-hoc two-sample *t*-tests at each age revealed that tutored birds were significantly more stereotyped than non-tutored at P65 (*t*(10) = 2.65, *p* = 0.024) but not at P90 (*t*(14) = 0.76, *p* = 0.46). The P90 condition difference remained non-significant, even after excluding the birds who did not sing at P65 (*t*(10) = 1.71, *p* = 0.118). While this null result at P90 should be interpreted with caution based on the small sample size available for this contrast, it may possibly reflect the fact that these ages, P65 and P90, occur within different phases within the song learning process, i.e., the plastic song phase and song crystallization phase, respectively (Johnson et al., 2002).

#### Predicting song stereotypy using FC features

To examine whether the FC features identified in previous analyses predicted song stereotypy, we implemented backwards stepwise model selection, beginning with the following predictors: condition, age, average ICC of the significant cluster that distinguished condition, developmental change in ICC from the previous age to the current age of singing, average S2V FC of the significant cluster that distinguished condition, and developmental change in S2V FC from the previous age of singing to the current age of singing. Stepwise model selection proceeded to eliminate (1) condition (ΔAIC = 1.79) and (2) developmental change in S2V FC (ΔAIC = 1.67). Further model reduction resulted in larger AIC values, and the final model is displayed in Table 3.1. This model explained 56.9% of total stereotypy variance (*Adjusted R^2^* = 0.49).

**Table 3.1.** Stereotypy Prediction.

| Predictor | Estimate | SE | B | T | p |
| --- | --- | --- | --- | --- | --- |
| Intercept | -20.60 | 11.86 | 0.00 | -1.74 | 0.096 |
| L NCM ICC | -7.50 | 3.87 | -0.40 | -1.94 | 0.065 |
| L NCM ICC Change | 9.47 | 2.25 | 0.75 | 4.21 | < 0.001 |
| L NCM - Cerebellum S2V FC | 53.41 | 17.71 | 0.52 | 3.02 | 0.006 |
| Age | 0.29 | 0.15 | 0.29 | 1.92 | 0.067 |

Note. $DfE = 23$ , $Rsquared = 0.569$ , Adjusted $Rsquared = 0.494$

We found that a larger change in L NCM ICC from the previous to current singing time point (i.e., P65 ICC minus P45 ICC, and P90 ICC minus P65 ICC) was a strong positive predictor of song stereotypy at the current age, at both P65 and P90 (*β* = 0.75, *t*(23) = 4.21, *p* < 0.001). Similarly, the average FC between the L NCM ICC cluster and the L Cerebellum S2V cluster positively predicted stereotypy (*β* = 0.52, *t*(23) = 3.02, *p* = 0.006). Age marginally positively predicted stereotypy (*β* = 0.29, *t*(23) = 1.92, *p* = 0.067). Finally, the effect of average L NCM ICC was non-significant and negative (*β* = −0.40, *t*(23) = −1.94, *p* = 0.065). Notably, this non-significant *negative* effect likely resulted from a statistical suppression effect (Tu, Gunnell, & Gilthorpe, 2008) and should not be interpreted as a true negative association. Specifically, the bivariate association between average L NCM ICC and stereotypy was positive (*β* = 0.35, *t*(26) = 1.90, *p* = 0.069), becoming negative only after conditioning on two moderately correlated covariates, L NCM ICC change (*r*(26) = 0.61, *p* < 0.001) and S2V FC (*r*(26) = 0.47, *p* = 0.011). The association of L NCM ICC and ICC change with stereotypy is visually depicted in Figure 6.

**Figure 6.**
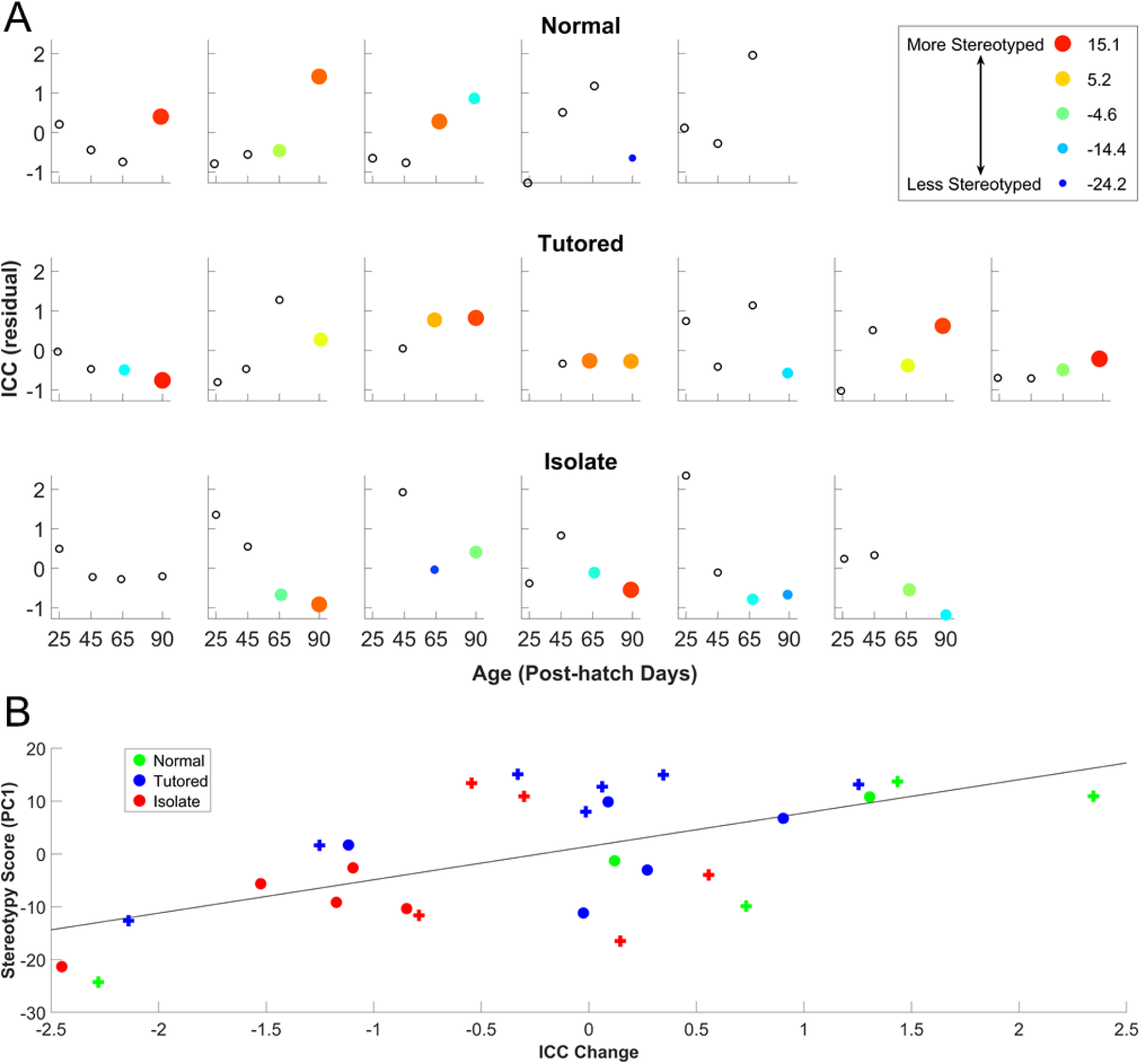
The association between developmental trends in ICC and stereotypy. **A:** Scatter plots for each individual bird showing the developmental trajectory of cluster ICC from P25 to P90. Hollow data points denote a lack of stereotypy data for a given age (*x*-axis), whereas both the color and size of filled data points denote the level of stereotypy in units of the first principal component scores. ICC scores (*y*-axis) are residuals from a regression controlling for the effects of body temperature, isoflurane dose, and head coil. **B:** A scatter plot showing the association between change in ICC from the previous age and stereotypy across all birds. An OLS regression line is displayed (*R^2^* = 0.37). Ages P65 (o) and P90 (+) are denoted with distinct markers.

#### FC mediates stereotypy differences between rearing conditions

Given that ICC change from the previous age was the strongest predictor of stereotypy, and rearing condition was no longer a significant predictor when controlling for this factor and other covariates, we conducted a mediation analysis to determine whether ICC change statistically mediated rearing condition-related differences in stereotypy. We found that ICC change did significantly mediate the rearing condition effect (*ACME* = 0.43, *CI*: [0.08, 0.98], *p* = 0.022). The portion of the rearing condition effect *not* mediated by ICC change was non-significant (*ADE* = 0.31, *CI*: [-0.43, 1.06], *p* = 0.388), whereas the total effect (i.e., a combination of the *ACME* and *ADE*) was significant (*TE* = 0.75, *CI*: [-0.01, 1.41], *p* = 0.046). ICC change mediated approximately 58.1% of the total effect (*CI*: [11.3%, 544%], *p* = 0.061). A mediation path diagram and the bootstrap distribution of the *ACME* are displayed in Figure 7.

**Figure 7.**
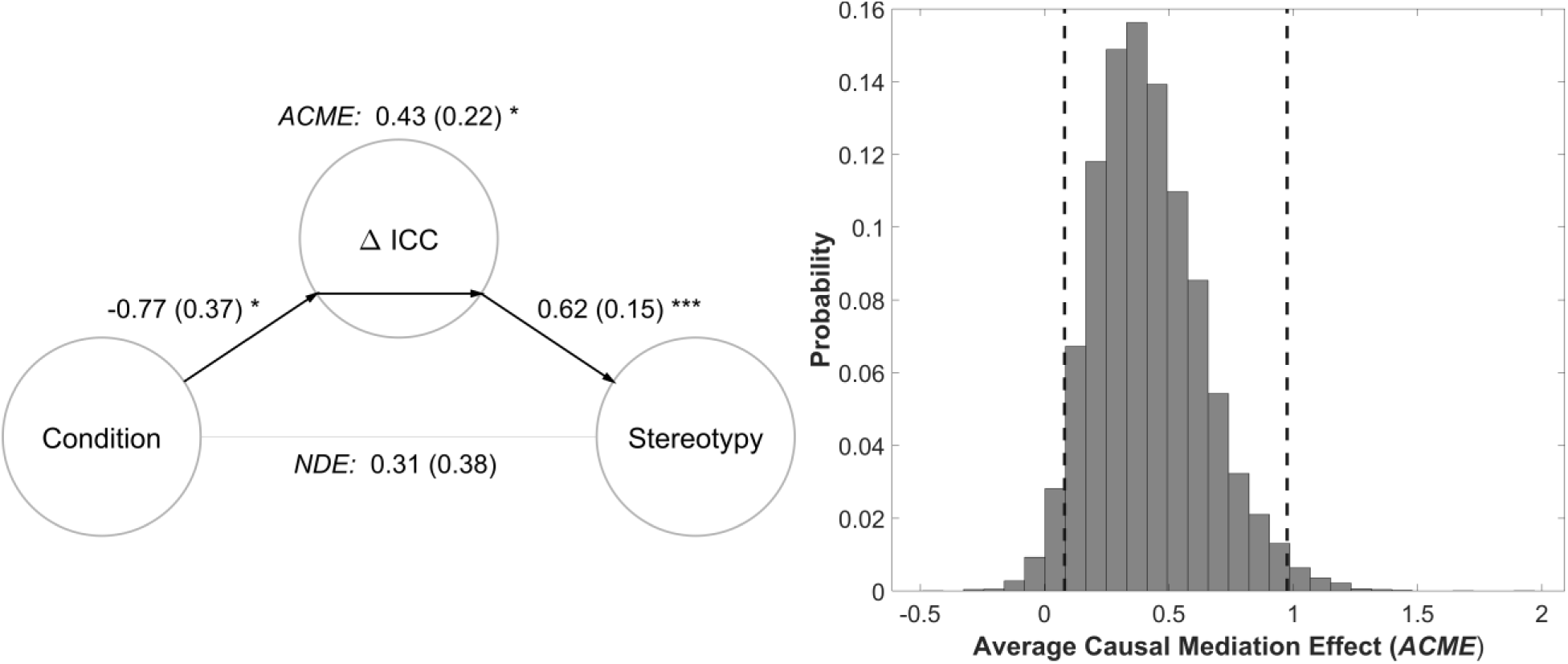
Mediation of rearing condition-related differences in stereotypy. **Left:** Path model for the mediation of rearing condition-related differences in stereotypy by change in left NCM ICC from the previous age. Coefficient estimates are reported with standard errors in parentheses. \**p* < 0.01, \*\**p* < 0.005, \*\*\**p* < 0.001. The left panel was created using custom Matlab scripts and the Mediation Toolbox (Wager, Davidson, Hughes, Lindquist, & Ochsner, 2008). **Right:** A probability scale histogram of the bootstrap distribution for the indirect effect, also known as the average causal mediation effect (*ACME*). Dashed lines demarcate the 95% confidence interval of the mediation effect, which notably does not overlap zero.

## Discussion

The current study identified FC signatures of social tutor experience across the CP for song learning in zebra finches. Data-driven whole-brain analyses identified a cluster centered within left NCM in which ICC decreased across the CP for Isolates but increased or was stable for Normal and Tutored birds. S2V analyses revealed that the FC between the left NCM cluster and a cluster centered within the left dorsal cerebellum exhibited parallel developmental trends. Interestingly, both the change in left NCM ICC and the magnitude of FC between the left NCM and dorsal cerebellum clusters strongly predicted the juvenile zebra finches’ song stereotypy at P65 and P90, in the middle and towards the end of the sensorimotor learning period. Additionally, changes in left NCM ICC from the previous time point statistically mediated non-tutored birds’ lower levels of stereotypy.

Together, these findings suggest that the developmental trajectory of left NCM FC may be highly dependent upon social tutor experience during the CP, consistent with previous reports that NCM undergoes experience-dependent neural plasticity in support of tutor song memorization (for reviews, see Bolhuis & Moorman, 2015; London, 2017). Additionally, the association between left NCM ICC and the subsequent emergence of song stereotypy during sensorimotor learning suggests that the functional connections of the auditory forebrain may play an important role in successful sensorimotor learning (Akutagawa & Konishi, 2010; Bauer et al., 2008; Fortune & Margoliash, 1995; Shaevitz & Theunissen, 2007; Vates et al., 1996; Yanagihara & Yazaki-Sugiyama, 2016).

The observation that left NCM ICC depends upon tutor experience during the CP is consistent with a large and growing body of evidence indicating that NCM undergoes tutor experience-dependent plasticity in support of tutor song memorization (Ahmadiantehrani & London, 2017; Kelly, Ahmadiantehrani, Blattler, & London, 2018; London & Clayton, 2008; Yanagihara & Yazaki-Sugiyama, 2016). The current results extend these findings to the systems level using rs-fMRI. Moreover, they indicate that, not only does *local* tutor experience-dependent plasticity occur within NCM itself, but tutor experience also appears to support the development of NCM’s centrality or importance within broader functional networks. Specifically, the developmental decline in ICC observed within the left NCM of Isolates suggests that tutor experience during the CP may be critical for preserving or strengthening functional connections between NCM and other regions. If tutor experience does not occur, then NCM FC may weaken with relative disuse.

Interestingly, some evidence suggests that NCM may be functionally left lateralized for tutor song sensory learning (Chirathivat, Raja, & Gobes, 2015; Moorman et al., 2012; Moorman & Nicol, 2015; Olson, Maeda, & Gobes, 2016; cf. Phan & Vicario, 2010; Voss et al., 2007). For instance, Moorman et al. (2012) found that NCM IEG *ZENK* expression was left lateralized specifically in response to tutor song playback during the sensory learning period, but not for the playback of novel song or silence (Moorman et al., 2012). Moreover, the degree of NCM left lateralization was strongly correlated with the extent of tutor song copying (Moorman et al., 2012). A similar result was reported for spontaneous NCM *ZENK* expression in juveniles during sleep: good learners (i.e., superior tutor song copying) showed significantly more left lateralization for NCM activity, whereas poor learners tended to be more right lateralized (Moorman, Gobes, van de Kamp, Zandbergen, & Bolhuis, 2015). In the current study, we identified a cluster confined within the left hemisphere, wherein ICC was developmentally dependent on tutor experience. However, a follow-up analysis revealed that a homotopic right NCM cluster also showed a parallel rearing condition-related developmental difference in ICC. The effect size in the right NCM cluster was numerically smaller than that observed in left NCM, but these effects were not significantly different (*p* = 0.224). It is possible that the current study was simply insufficiently powered to detect a small lateralization effect. However, this result also seems to indicate that the condition-related developmental difference observed in NCM was, at least to some extent, bilateral in nature. The cluster detected did not extend into the right hemisphere possibly due to the use of a stringent CDT, which is necessary for valid statistical inference with the cluster-extent threshold method (e.g., Woo, Krishnan, & Wager, 2014). Notably, the lack of a lateralization effect also seems consistent with our previous observation of strong homotopic FC between the left and right auditory forebrain (Layden et al., 2019).

In addition to left NCM, the present results also highlighted the potential involvement of a novel cluster encompassing part of the left dorsal cerebellum. Although the cerebellum has a widely recognized, if incompletely understood, role in fine motor control (for review, see Manto et al., 2012), its potential contribution to song learning and/or production has received relatively little attention. In contrast, a large and growing body of evidence suggests that the cerebellum plays important roles in various aspects of human language (for reviews, see Hertrich, Mathiak, & Ackermann, 2016; Pleger & Timmann, 2018; Ziegler & Ackermann, 2017), including linguistic prediction (D’Mello, Turkeltaub, & Stoodley, 2017; Lesage, Hansen, & Miall, 2017; Lesage, Morgan, Olson, Meyer, & Miall, 2012; Moberget, Gullesen, Andersson, Ivry, & Endestad, 2014), verbal working memory (Chen & Desmond, 2005; Kirschen, Chen, Schraedley-Desmond, & Desmond, 2005; Ravizza et al., 2006), and word generation (Arasanz, Staines, Roy, & Schweizer, 2012). Recently, a link was found between the songbird anterior forebrain pathway (AFP) and the cerebellum (Perkel, 2004). Previously, it was known that Area X (basal ganglia) of the AFP projects to the dorsolateral nucleus of the anterior thalamus (DLM). However, it was only recently discovered that Area X also receives a projection from DLM, as well as from an adjacent region that provides cerebellar input (Hamaide et al., 2018; Nicholson, Roberts, & Sober, 2018). Although this connection does not provide a direct structural connectivity pathway between NCM and the cerebellum, it does at least establish a structural link between the cerebellum and the song network. Given the parallels between human speech acquisition and zebra finch song learning, this link, and potentially other yet undiscovered links, between the cerebellum and song network may merit further investigation for potential roles in song learning or production.

Although our results suggest that left NCM ICC is linked to the emergence of song stereotypy, we were unable to determine whether left NCM ICC was also related to the extent of tutor song copying. Tutor song similarity data was unavailable for Isolates, given that they were not tutored, but it was also not known which adult males served as tutors for the aviary-reared Normal condition birds. Future studies might utilize a larger sample of Tutored condition birds to examine whether left NCM FC is also associated with the extent of tutor song copying, in addition to song stereotypy.

## Conclusion

Overall, our results suggest that tutor experience during the CP for tutor song memorization strongly modulates the development and maintenance of left NCM FC, consistent with prior indications that NCM is critical for tutor song memorization. Moreover, left NCM ICC strongly predicted the emergence of song stereotypy during the sensorimotor learning period. Future investigations are merited to determine whether left NCM FC similarly predicts the extent of tutor song copying, and we hope that the current study lays the groundwork for such investigations. Importantly, the use of rs-fMRI allowed us to obtain repeated measures of *in vivo* whole-brain FC from the same intact animals at multiple developmental time points, a feat which may be difficult to achieve using more invasive techniques. We therefore believe that the current study underlines the utility of rs-fMRI for conducting comparative studies to investigate the neural mechanisms that support the developmental learning of complex behaviors.

